# Integrative Network Biology Framework Elucidates Molecular Mechanisms of SARS-CoV-2 Pathogenesis

**DOI:** 10.1101/2020.04.09.033910

**Authors:** Nilesh Kumar, Bharat Mishra, Adeel Mehmood, Mohammad Athar, M. Shahid Mukhtar

## Abstract

COVID-19 (Coronavirus disease 2019) is a respiratory illness caused by severe acute respiratory syndrome coronavirus 2 (SARS-CoV-2). While the pathophysiology of this deadly virus is complex and largely unknown, we employ a network biology-fueled approach and integrated multiomics data pertaining to lung epithelial cells-specific coexpression network and human interactome to generate <u>C</u>alu-3-specific human-<u>S</u>ARS-CoV-2 <u>I</u>nteractome (CSI). Topological clustering and pathway enrichment analysis show that SARS-CoV-2 target central nodes of host-viral network that participate in core functional pathways. Network centrality analyses discover 28 high-value SARS-CoV-2 targets, which are possibly involved in viral entry, proliferation and survival to establish infection and facilitate disease progression. Our probabilistic modeling framework elucidates critical regulatory circuitry and molecular events pertinent to COVID-19, particularly the host modifying responses and cytokine storm. Overall, our network centric analyses reveal novel molecular components, uncover structural and functional modules, and provide molecular insights into SARS-CoV-2 pathogenicity.

## Introduction

From the epicenter of the COVID-19 (Coronavirus disease 2019) outbreak in China, the disease has spread globally in 185 countries/territories with over 1.4 million confirmed cases and almost 87,000 fatalities as of April 07, 2020, and the World Health Organization (WHO) warned that the pandemic is accelerating worldwide^1, 2^. Apart from the human tragedy, COVID-19 has a growing detrimental impact on the global economy and will likely cause trillions in financial losses worldwide in 2020 alone. COVID-19 is an infectious respiratory illness caused by a highly contagious and pathogenic SARS-CoV-2 (severe acute respiratory syndrome coronavirus 2). This single-stranded RNA virus belongs to the family *Coronaviridae* and is closely related to another human coronavirus SARS-CoV with 89.1% nucleotide similarity^2, 3^. SARS-CoV and another human coronavirus MERS-CoV (Middle East Respiratory Syndrome-CoV) caused two previous global epidemics in 2003 and 2012, respectively, both characterized by high fatality rates^2, 3^. These coronaviruses mainly spread from a contagious individual to a healthy person through respiratory droplets derived from an infected person’s cough or sneeze, and from direct contact with contaminated surfaces or objects, where the virus can maintain its viability for period ranging from hours to days^2, 3^. Unlike other coronaviruses, SARS-CoV-2 transmits more efficiently and sustainably in the community according to Center for Disease Control (CDC)^4^. While majority of the patients infected with SARS-CoV-2 develop a mild to moderate self-resolving respiratory illness, infants and older adults (≥ 65 years) as well as patients with preexisting medical conditions such as cardiovascular disease, diabetes, chronic respiratory disease, renal dysfunction, obesity and cancer are more vulnerable^2, 3^. The pathophysiology of SARS-CoV-2 is complex and largely unknown but is associated with an extensive immune reaction referred to as ‘cytokine storm’ triggered by the excessive production of interleukin 1 beta (IL-1β), interleukin 6 (IL-6) and others. The cytokine release syndrome leads to extensive tissue damage and multiple organ failure^2^. While no vaccine or antiviral drugs are currently available to prevent or treat COVID-19, identifying molecular targets of the virus could help uncover effective treatment. Towards this, generation of a human-SARS-CoV-2 interactome, integration of virus-related transcriptome to interactome, discovery of disease-related structural and functional modules, and dynamic transcriptional modeling will provide insights into the virulent mechanisms of this deadly virus.

Networks encompass a set of nodes and edges, also referred as vertices and links, respectively. Nodes are systems components, whereas edges represent the interactions or relationships among the nodes^5, 6^. In biological systems, genes and their products perform their functions by interacting with other molecular components within the cell. For instance, proteins directly or indirectly interact with each other under both steady-state and different stress conditions to form static and dynamic complexes, and participate in diverse signaling cascades, distinct cellular pathways and a wide spectrum of biological processes. Proteome scale maps of such protein-protein interactions are referred to as interactomes^5, 6^. Meanwhile, specialized pathogens including viruses, bacteria and eukaryotes employ a suite of pathogenic or virulent proteins, which interact with high value targets in host interactomes to extensively rewire the flow of information and cause disease^5, 7, 8, 9^. Therefore, analyzing the network architecture and deciphering the structural properties of host-pathogens interactomes may reveal novel components in virus pathogenicity. Such analysis indicates that diverse cellular networks are governed by universal laws and exhibit scale-free network topology, whose degree distribution follows a power law distribution with a few nodes harboring increased connectivity^5^. Given that diverse biological systems display similar network architecture and topology, several structural features and physical characteristics within a cellular network may act as indicators of important nodes^5^. These include degree (the number of edges of a node), and betweenness (the fraction of the shortest paths that include a node)^6, 10, 11^. Indeed, it has been shown that hubs (high degree nodes) and high betweenness nodes (bottlenecks) are targets of numerous human–viral, human–bacterial and other human diseases^5, 7, 8, 9, 12, 14, 15, 16, 17, 18, 19, 20, 21, 22^. In addition, host protein targets of diverse pathogens were demonstrated to be in close proximity (shortest path) with differentially expressed genes (DEGs)^23^. Specific to viral-host pathosystem, the network analysis of several interactomes including human T-cell lymphotropic viruses, Epstein-Barr virus, hepatitis C virus, influenza virus and human papillomavirus indicate that several of the above described topological features are associated with viral targets^12, 13, 14, 15, 16, 17, 18, 19, 20, 24, 25, 26, 27, 28^. Recently, in SARS-CoV human interactome, nodes corresponding to hubs and bottlenecks including respiratory chain complex I proteins were identified as targets of SARS-CoV. This system-wide analysis also identified several immunophilins as direct physical interacting partners of the CoV non-structural protein 1 (Nsp1)^12^. Importantly, using affinity-purification mass spectrometry (AP-MS), a proteome-scale mapping recently identified 332 SARS-CoV-2 Interacting Proteins (SIPs) in human^29^. This groundbreaking study paved new avenues to investigate novel therapeutic targets using a systems pharmacology approach. Undoubtedly, network biology presents a nextgeneration, integrative approach for drug repurposing that can predict individual or, more likely, combinatorial sets of drugs with high efficacy against SARS-CoV-2^30, 31^.

Here, we generated a comprehensive human-SARS-CoV-2 interactome encompassing 12,852 nodes and 84,100 edges. We also performed a weighted co-expression network analysis (WGCNA) in Cultured Human Airway Epithelial Cells (Calu-3) treated with SARS-CoV or MERS-CoV over time (22, 445 nodes, 10,649,854 edges). By integrating co-expression network with interactome, we obtained <u>C</u>alu-3-specific human – <u>S</u>ARS-CoV-2 <u>I</u>nteractome (CSI) containing 4,123 nodes and 14,650 edges. Network analysis indicates that the average degree, betweenness and information centrality of SIPs are enriched in CSI. Module-based functional pathway analyses discovered several disease-related clusters that are enriched in several signaling pathways and biological processes including eIF2 signaling/translation, inhibition of ARE-mediated mRNA degradation pathway, protein ubiquitination pathway, T cell receptor regulation of apoptosis, NER pathway, RNA degradation and retinoic acid-mediated apoptosis signaling pathway. Network topology analyses identified 28 high-value targets of SARS-CoV-2, which can form complexes with other highly influential nodes within CSI. These most important nodes are possibly involved in the viral entry, proliferation and survival in the host tissue as well as required to induce a conducive environment for viral sustenance and pathogenesis. Moreover, we incorporated transcriptome data of COVID-19 patients derived from bronchoalveolar lavage fluid (BALF) and peripheral blood mononuclear cells (PBMC) with our CSI data. Subsequently, we performed dynamic gene regulation modeling on the CSI nodes to decipher the intricate relationships between important transcription factors (TFs) and their target genes upon SARS-CoV-2 infection. Of particular of interest is the TF-regulatory relationships involved in host modifying processes such as protein translation, ubiquitination, and the cytokine storm. In summary, our integrative network topology analyses led us elucidate the underlying molecular mechanisms and pathways of SARS-CoV-2 pathogenesis.

## Results

### Integrated interactome-transcriptome analysis to generate <u>C</u>alu-3-specific human- <u>S</u>ARS-CoV-2 <u>I</u>nteractome (CSI)

It is likely that the outcome of SARS-CoV-2 infection can largely be determined by the interaction patterns of host proteins and viral factors. To build the human – SARS-CoV-2 interactome, we first assembled a comprehensive human interactome encompassing experimentally validated PPIs from STRING database^32^. Since the STRING database is not fully updated, we manually curated PPIs from four additional proteomes-scale interactome studies, *i.e.* Human Interactome I and II, BioPlex, QUBIC, and CoFrac (reviewed in^33^). This yielded us an experimentally validated high quality interactome containing 18,906 nodes and 444,633 edges (Fig. 1a). Subsequently, we compiled an exhaustive list of 394 host proteins interacting with the novel human coronavirus that was referred to as <u>S</u>ARS-CoV-2 <u>I</u>nteracting <u>P</u>roteins (SIPs) (Supplementary Data 1). This comprises 332 human proteins associated with the peptides of SARS-CoV-2^29^, whereas the remaining 62 host proteins interact with the viral factors of other human coronaviruses including SARS-CoV and MERS-CoV^34^, which could also be of significance in understanding the molecular pathogenesis of SARS-CoV-2. By querying these 394 SIPs in the human interactome, we generated a subnetwork of 12,852 nodes and 84,100 edges that covers first and second neighbors of 373 SIPs (Fig. 1a). Given that the SIPs-derived PPI subnetwork may not operate in all spatial or temporal conditions, coronavirus-specific co-expression data is used to filter the interactions in the context of COVID-19. It is important to note that no exceptionally high-resolution SARS-CoV-2 transcriptome was available at the time of analysis (details below). Therefore, we took advantage of extensive temporal expression data available for SARS-CoV and MERS-CoV (Fig. 1b). Towards this, we performed a weighted coexpression network analysis (WGCNA) in Human Airway Epithelial Cells (Calu-3) treated with SARS-CoV and MERS-CoV over time *in vitro* in culture. This analysis yielded a comprehensive co-expression network with 22,445 nodes and 10,649,854 edges (Fig. 1b). By integrating this Calu-3 co-expression network with SIPs-derived PPI subnetwork, we generated <u>C</u>alu-3-specific human-<u>S</u>ARS-CoV-2 <u>I</u>nteractome (CSI) that contains 214 SIPs interacting with their first and second neighbors make a network of 4,123 nodes and 14,650 edges (Fig. 1c, Supplementary Data 1). We showed that CSI follows a power law degree distribution with a few nodes harboring increased connectivity, and thus exhibits properties of a scale-free network (*r^2^* = 0.91; (Fig. 1d, Supplementary Data 1), similar to the previously generated other human-viral interactomes^12, 13, 14, 15, 16, 17, 18, 19, 20, 24, 25, 26, 27, 28^. Taken together, we constructed a robust, high quality CSI that was further utilized for network-aided architectural and functional pathway analyses.

**Figure 1:**
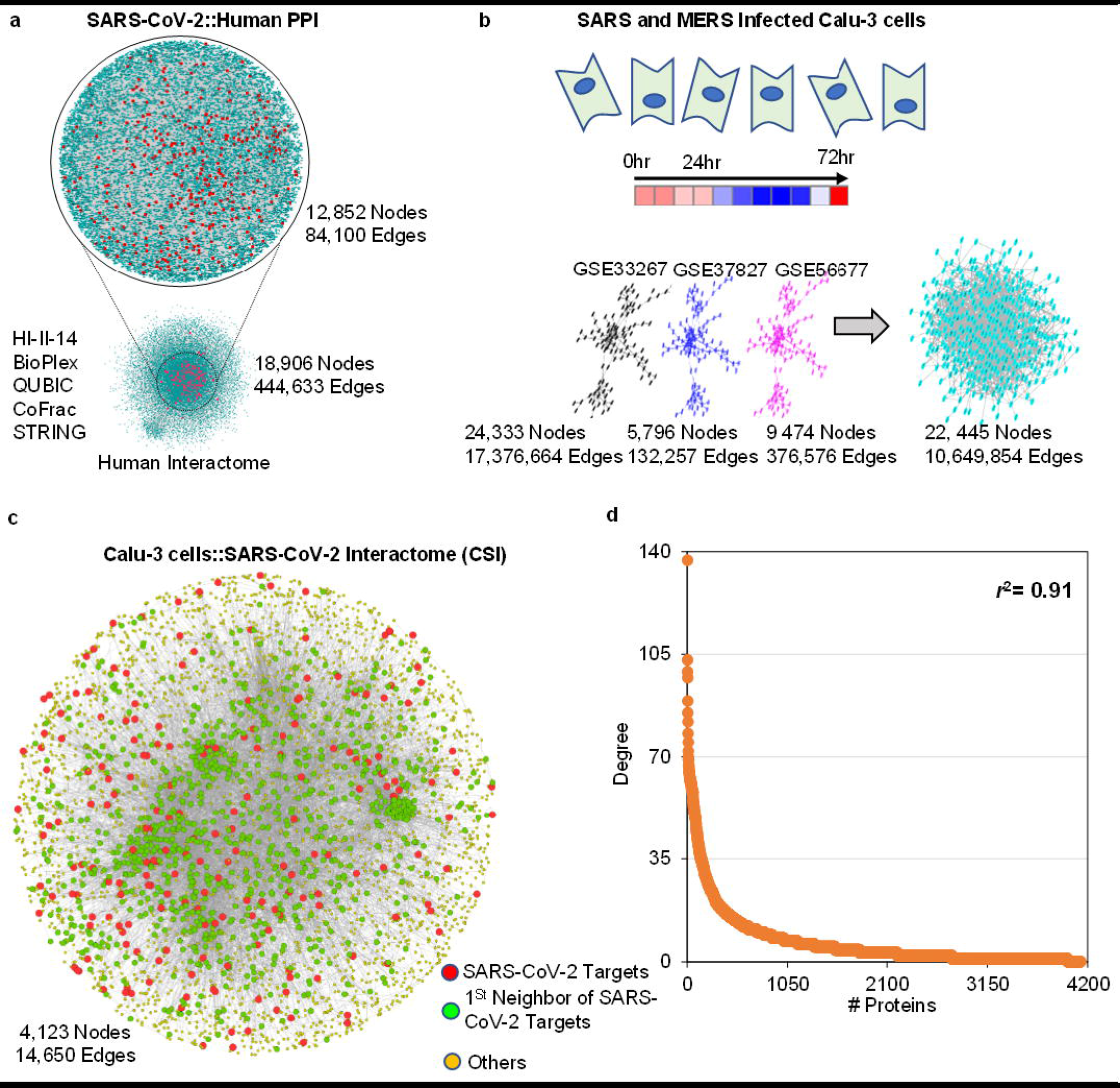
Integrative multi-omics analysis identified Calu-3-specific human-SARS-CoV-2 Interactome (CSI). **a** Human interactomes (HI-II-14, BioPlex, QUBIC, CoFrac, and STRING) connections and 373 SARS-CoV-2 Interacting Proteins (SIPs) were used to extract the “SARS-CoV-2::Human PPI” (12,852 Nodes and 84,100 Edges) including all possible interactions. **b** Weighted gene co-expression network (WGCNA) construction of SARS and MERS Infected Calu-3 cells gene expressions profiles from NCBI GEO datasets. The merged co-expression network has 22,445 Nodes and 10,649,854 Edges. **c** Calu-3-specific human-SARS-CoV-2 Interactome (CSI) with 4,123 Nodes and 14,650 Edges (Red: 214 SIPs, Green: 1^st^ Neighbor of SARS-CoV-2 Interacting Proteins (SIPs), Yellow: other proteins). **d** Degree of CSI nodes displays power law (*r^2^* =0.91) distribution and follow scale free property.

### Network topology and module-based functional analyses reveal that SARS-CoV-2 targets core signaling pathways of the host network

From a network biology standpoint, a viral infection as well as other pathogen attacks can be viewed as a set of strategic perturbations, at least in part, within the core components of the host interactome^24, 25, 26^. Since such central nodes correspond to proteins that exhibit increased connectivity and/or central positions within a network, we addressed a question whether SARS-CoV-2 also attacks such important nodes within CSI. Towards this, we calculated the average degree (number of connections), betweenness (the fraction of all shortest paths that include a node within a network), load centrality (the fraction of all shortest paths that pass through a node), information centrality (the harmonic mean of all the information measures for a node in a connected network) and PageRank index (counting incoming and outgoing connections considering the weight of the edge) for SIPs, and compared them with their first and second neighbors. We demonstrated that these four topological features of SIPs were significantly higher than the other nodes within CSI (Fig. 2a, b, c and Supplementary Fig. 1a and b, Supplementary Data 2; *t*-test *P* < 0.0001). We also showed that SIPs were significantly enriched in CSI compared to the human interactome (Fig. 2d, Supplementary Data 2; hypergeometric *P* < 3.159E-51). These results indicate that SARS-CoV-2 targets core structural components of the human-viral interactome, and prompted another question as to whether CSI also activates common biological processes in response to viral infection. Since nodes within CSI not only form protein complexes with each other but also transcriptionally co-express, we reasoned that densely connected nodes within this network may participate in similar biological functions. Towards this, we investigated the underlying modular structures (protein clusters ≥ 5 nodes) in CSI followed by Ingenuity Pathway Analysis (IPA). This approach allowed us to identify 27 modules ranging from 5 to 66 nodes for the smallest and largest modules, respectively. Subsequently, we examined the biological processes, cellular pathways and signaling cascades that are modulated in the top 10 modules (Fig. 2e-k, Supplementary Data 2). Significantly enriched signaling pathways and biological processes included eIF2 signaling/translation representing protein translation control, inhibition of ARE-mediated mRNA degradation pathway, protein ubiquitination pathway, T cell receptor regulation of apoptosis, NER pathway, RNA degradation and retinoic acid-mediated apoptosis signaling pathway (-log(*P*-value) ≥ 2; Fig. 2 e-k, Supplementary Fig. 1c, Supplementary Data 2). Enrichment analysis of KEGG pathways can be utilized to ascertain signal transduction, as well as biochemical and metabolic pathways. The significantly enriched pathways (*P*-value ≤0.05) mainly included the infection with a number of viruses, oxidative phosphorylation, ER protein processing and apoptosis (Supplementary Fig. 1d, Supplementary Data 2). Finally, we performed a human phenotype ontology analysis that identifies phenotypic abnormalities encountered in human diseases. Significantly enriched terms included mitochondrial inheritance, hepatic necrosis, respiratory failure and abnormality of the common coagulation pathway (Supplementary Fig. 1e). Collectively, we showed that SARS-CoV-2 proteins interact with central nodes of CSI, and these proteins are implicated in core molecular and cellular pathways to establish infection and continue disease progress.

**Figure 2:**
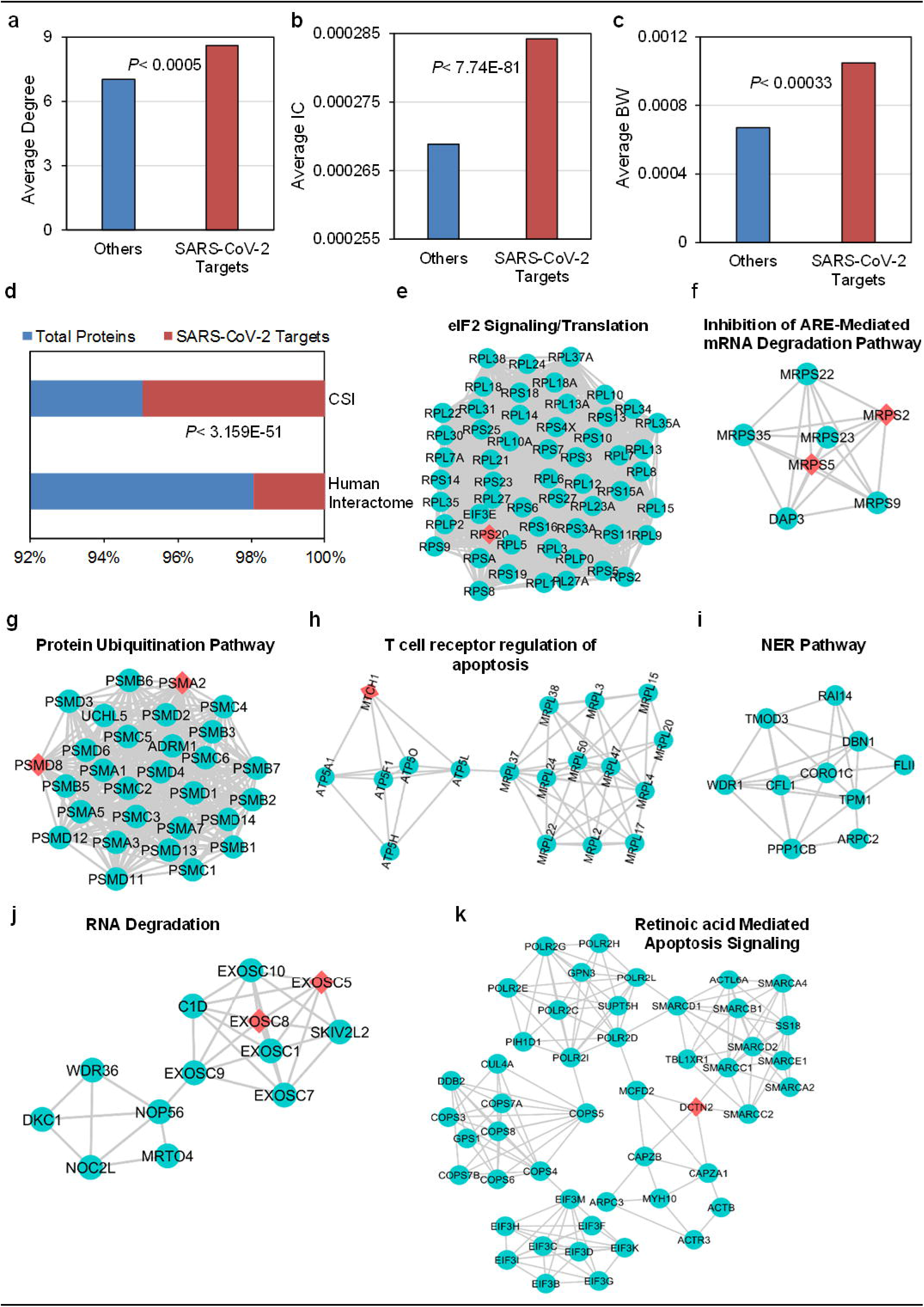
SARS-CoV-2 Interacting Proteins (SIPs) structural and functional properties in Calu-3-specific human-SARS-CoV-2 Interactome (CSI). **a** Average degree of SIPs (8.61) is significantly higher than other interacting proteins (7.03) in CSI network *(t-* test, *P* < 0.0005). **b** Average information centrality (IC) of SIPs (0.000284) is significantly heighted compared to remaining proteins (0.000269) in CSI network (*t*-test, *P* < 7.73E-81). **c** SIPs exhibit significantly increased average betweenness centrality (BW; 0.00105) compared to other interacting proteins (0.00067) in CSI network (*t*- test, *P* < 0.00033). **d** SIPs are significantly enriched in CSI network than the human interactome (hypergeometric test, *P* < 3.159E-51). **e-k** CSI subnetworks of highly clustered modules obtained with the application of MCODE Cytoscape app and *K*-means clustering. The significant functional annotation was done by Ingenuity Pathway Analysis (IPA) (Red= SARS-CoV-2 targets, Olive Blue= CSI nodes).

### Network topology framework identifies most influential nodes in CSI

Human-viral interactome landscapes of several viruses have previously shown that viral proteins interact with nodes corresponding to high degree (hubs) and high betweenness (bottlenecks), and such structural features have been previously used to predict viral targets^12, 13, 14, 15, 16, 17, 18, 19, 20, 24, 25, 26, 27, 28^. In addition to hubs and bottlenecks, PageRank algorithm was also effectively used to identify viral targets^27^. Moreover, these physical characteristics can also be used to prioritize the most influential genes in CSI for biological relevance and drug target discovery. Here, we used nine different centrality indices to identify the most influential nodes referred to as <u>C</u>SI <u>S</u>ignificant <u>P</u>roteins (CSPs). This includes the above described degree, betweenness, information centrality, PageRank index and load centrality as well as additional features such as eigenvector centrality (a measure of the influence of a node in a network), closeness centrality (reciprocal of the sum of the length of the shortest paths between the node and network), harmonic centrality (reverses the sum and reciprocal operations of closeness centrality and weighted *k*-shell decomposition) (an edge weighting method based on adding the degree of two nodes in network partition). While weighted *k*-shell decomposition analysis was recently performed to increase the predictability of host targets of bacterial pathogens^35^, we showed that the top 5% of nodes reside in the inner layers of CSI (Fig. 3a, Supplementary Data 2). For other centrality measures, we also maintained a stringent threshold of top 5% to be considered as a highly influential node or CSP. Evidently, we can expect overlapping topological features for the same set of nodes. Noticeably, we observed a strong positive correlation between information centrality and degree (Fig. 3b; *r^2^* = 0.9), betweenness and degree (Fig. 3c; *r^2^* = 0.51) and PageRank and degree (Fig. 3d; *r^2^* = 0.84, Supplementary Data 2).

**Figure 3:**
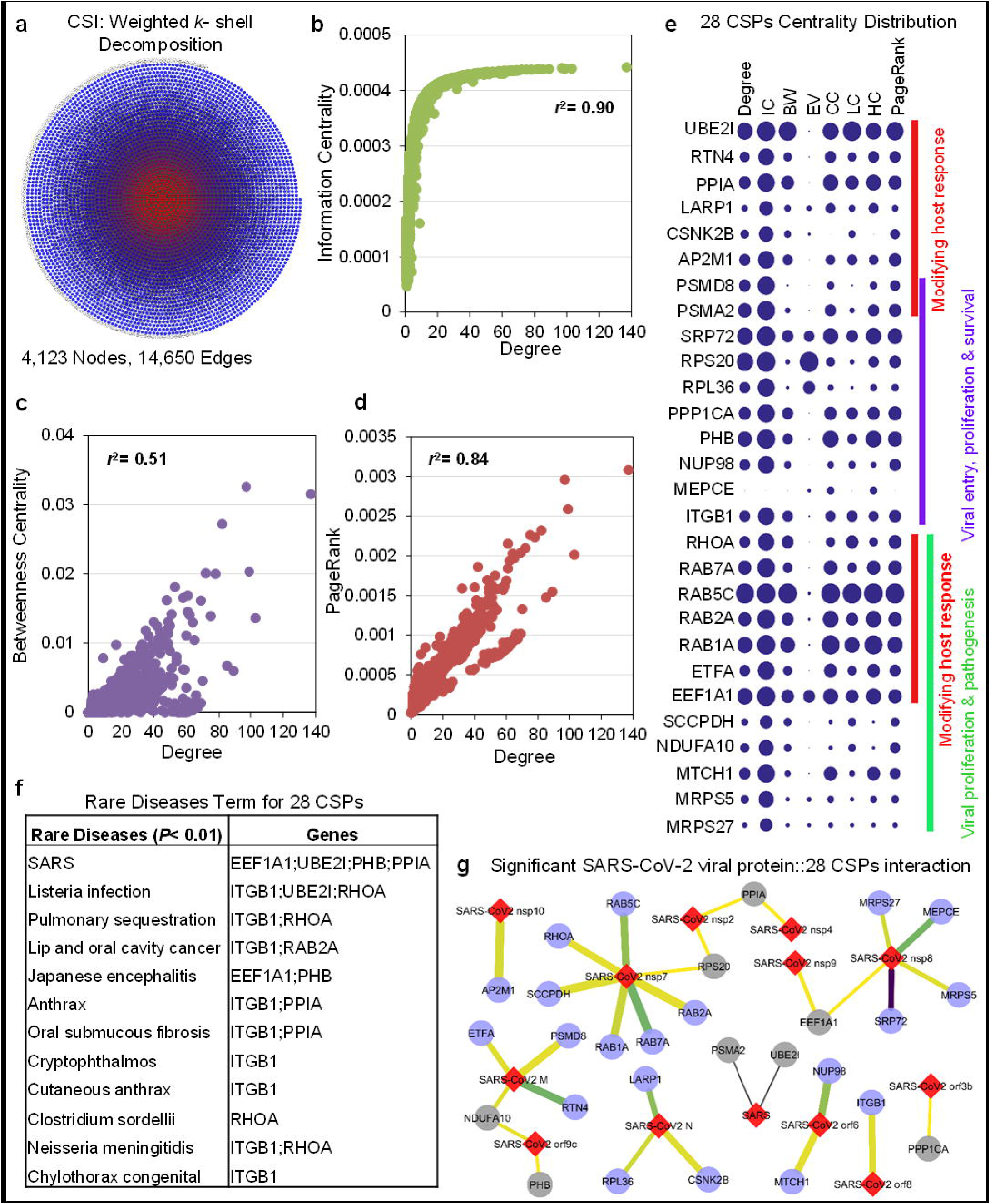
Identification of most influential nodes in Calu-3-specific human-SARS-CoV-2 Interactome (CSI) using Network biology framework. **a** Weighted *k*-shell decomposition identifies inner and peripheral layers of CSI. 33% of all shells are considered as inner layers. Top 5% of inner layer proteins are considered as significant (Red= Inner layer, Blue= peripheral layer). **b-d**. Correlation between information centrality and degree (**b**, *r^2^* = 0.9), betweenness and degree (**c**, *r*^2^ = 0.51) and PageRank and degree (**d**, *r*^2^ = 0.84). **e** 28 CSI Significant Proteins (CSPs) that exhibit more than one high significant centrality measures (Degree, information centrality (IC), betweenness centrality (BW), eigenvector centrality (EV), closeness centrality (CC), load centrality (LC), harmonic centrality (HC), and PageRank). The size of blue spot determines the significant central node in a centrality indices. **f** Enrichr identified significantly enriched rare disease enriched CSPs in SARS, listeria infection, and pulmonary sequestration (*P*-value <0.0001). **g** Network representation of significant SARS-CoV-2 viral protein interaction with 28 CSPs (Nodes: Red= viral proteins, Blue= CSPs significantly targeted by viral protein, grey= CSPs with insignificant viral protein interaction; edge width= MIST score, edge color= AvgSpec).

Collectively, we identified 28 CSPs that exhibit more than one high centrality measure (Fig. 3e, Supplementary Data 3). For instance, EEF1A1 that has previously been implicated in SARS was enriched in all the centrality measures tested in our study (Fig. 3f, Supplementary Data 3). In addition, UBE2I, PPIA, and PHB were also associated with SARS and were enriched in more than five centrality measures (Fig. 3f, Supplementary Data 3). We categorized these 28 CSPs into three major groups based on their potential roles in COVID-19. While we expect some, if not all, of these proteins to have more than one function, the group-1 CSPs might be largely relevant to modifying host response following SARS-CoV-2 infection (Fig. 3e). Moreover, the proteins in the other two groups might be involved in viral entry, proliferation, survival and pathogenesis as well as cytokine storm (Fig. 3e; see details in discussion). Furthermore, we found that these 28 CSPs are targets of some of the well-known SARS-CoV-2 viral proteins. SARS-CoV-2 nsp7 targets most of the CSPs *(i.e.* seven in total), SARS-CoV2 nsp8 targets five CSPs, and SARS-CoV-2 M has four CSPs targets, while other SARS-CoV-2 nsps’ (2,4,10) and SARS-CoV-2 orfs’ (3b, 6,8,9c) possess relatively fewer targets. Intriguingly, three of our CSPs (PPIA, RPS20, and NDUFA10) are targets of more than one SARS-CoV2 protein (Fig 3g, Supplementary Fig. 2), while PHB is the target of several viral proteins tested as bait at low threshold. It is also important to note that PHB is also targeted by viral proteins of SARS-CoV^34^. These data support previous findings that an individual viral factor can target multiple host nodes and several viral proteins can interact with the same host protein^12, 13, 14, 15, 16, 17, 18, 19, 20, 24, 25, 26, 27, 28^. Collectively, these data strengthen our notion that centrality measures can be an effective method to predict highly influential nodes, leading us to discover 28 such CSPs.

### Dynamic gene regulation modeling elucidates core transcriptional circuitry and regulatory signatures pertinent to SARS-CoV-2 infection

To further understand the biological characteristics, regulatory relationships and molecular events associated with the nodes in CSI, we incorporated transcriptome data of COVID-19 patients derived from bronchoalveolar lavage fluid (BALF) and peripheral blood mononuclear cells (PBMC) with our CSI data^36^. Overall, SARS-CoV-2 infection exhibited largely different transcriptional signatures for BALF and PBMC^36^. We identified a set of 228 and 215 differentially expressed genes (DEGs) in BALF and PBMC, respectively (*p* ≤ 0.05, FC ≥2.0, Fig. 4a, b, Supplementary Data 4). Thus, CSI constitutes over 25% of transcriptomes pertaining to both BALF and PBMC. Intriguingly, in BALF, we observed that the upregulated cluster A is enriched with eIF2 signaling/translation pathway, while the two down-regulated clusters (B and C) are enriched in retinoic acid-mediated apoptosis signaling pathway (Fig. 4a). Conversely, one major cluster that is significantly upregulated in PBMC is enriched in T cell receptor regulation of apoptosis and protein ubiquitination pathway (Fig. 4b). These data further support the notion that significantly enriched protein modules in CSI are involved in SARS-CoV-2 pathogenesis.

**Figure 4:**
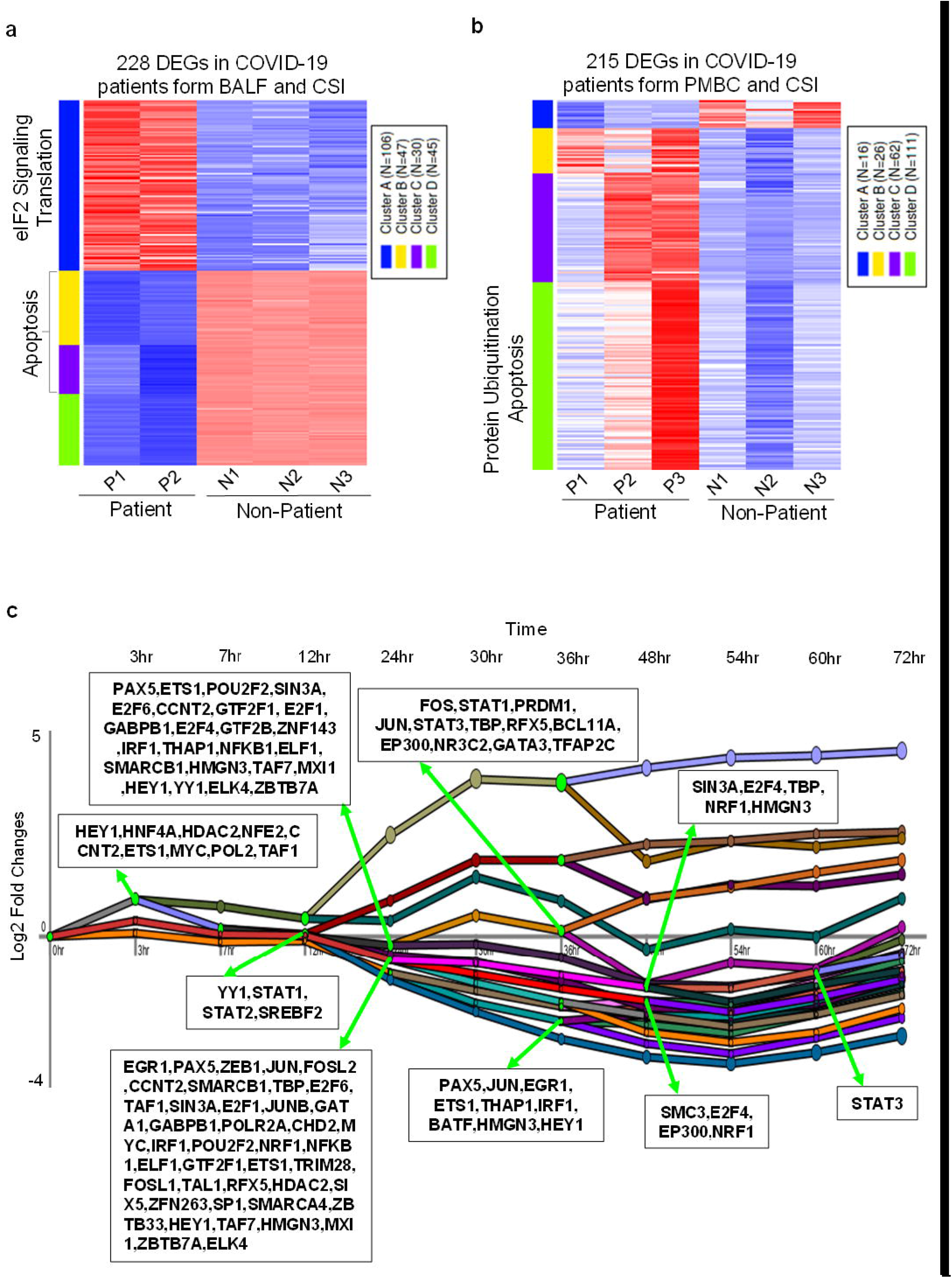
Dynamic gene regulation modeling of transcriptional signatures pertinent to SARS-CoV-2 infection. **a** Heatmap of 228 Differentially expressed genes (DEGs) in COVID-19 patients derived bronchoalveolar lavage fluid (BALF) transcriptome and <u>C</u>alu-3-specific human-<u>S</u>ARS-CoV-2 <u>I</u>nteractome (CSI). The heatmap was clustered based on *k*-mean with cluster with maximum genes are enriched in eIF2 Signaling/Translation pathways. Two out of three remaining clusters are enriched in Apoptosis. P and N denote patients and controls, respectively. **b** Heatmap of 215 DEGs common between transcriptome of peripheral blood mononuclear cells (PMBC) derived from P (patients) and N (controls) and Calu-3-specific human-SARS-CoV-2 Interactome (CSI). The heatmap was clustered based on *k*-mean with cluster with maximum genes are enriched in protein ubiquitination and apoptosis. **e** Dynamic regulatory event mining of 4,952 cumulative DEGs in SARS-CoV across 72 hours of infection reconstructed by incorporating static protein-DNA interaction data with time series (GSE33267). The regulators only expressed in BALF and PBMC transcriptomes are highlighted. Significant regulators (TFs) control the regulation dynamics *(P* < 0.05). Major bifurcation of pathways occurs at 24-hour with a total of 46 TFs involved in dynamic modulation.

To reveal the regulatory circuitry and molecular events pertinent to SARS-CoV-2 infection, we performed probabilistic modeling using iDREM (interactive Dynamic Regulatory Events Miner) framework that incorporates protein-DNA interaction data with transcriptomics^37^. Given that iDREM requires time-course transcriptional profiling data, and *in vivo* or *in vitro* temporal SARS-CoV-2 transcriptome data is currently lacking, we made use of a high-resolution temporal SARS-CoV dataset (11 time points)^38^. However, we only focused on those upstream transcriptional factors (TFs) and downstream target genes that were also present in BALF and PBMC, which allowed us to mimic SARS-CoV-2-mediated dynamic regulatory networks. This dynamic regulatory modeling identified several bifurcation points, where a set of TFs regulates their potential co-expressed and downstream target genes (Fig. 4c, Supplementary Fig. 3, Supplementary Data 4). Based on the expression trajectories and path expression patterns, we identified a total of 84 and 94 significant regulators that were expressed in BALF and PBMC COVID-19 patients’ samples, respectively (P < 0.05, Supplementary Data 4). Among them, we observed the first major wave of differential regulation and activation of TFs at 12-hour post infection. At this bifurcation transcriptional event, we found a set of 4 TFs (YY1, STAT1, STAT2, and SREBF2), which were also expressed in BALF transcriptome. The next major bifurcation occurred at 24-hour post infection, comprising 39 and 43 TFs expressed in BALF and PBMC, respectively (Supplementary Data 4). While we found similar sets of target genes regulated by diverse sets of TFs at different stages of infection, we also discovered multiple combinations of TFs regulating similar sets of downstream genes (Fig. 4c). This reflects the intricate nature of dynamic regulatory relationships between TFs and their targets.

Next, we primarily focused on four major pathways/signaling events, *i.e.* cytokine storm, eIF2 signaling/translation, protein ubiquitination pathway and T cell receptor regulation of apoptosis. In the first example of cytokine storm, we identified a total of 10 TFs (STAT2, SUZ12, JUN, STAT1, MEF2A, RAD21, STAT3, BCL11A, NFE2, and BATF) expressed in BALF and PBMC COVID-19 patients’ samples (Supplementary Data 4). Predominantly, we found that two TFs (STAT1 and STAT2) and one master regulator (JUN) are early transcriptional players activated at 12- and 24-hour post infection. In particular, we found CSI genes CXCL1 and TNFAIP3 co-regulated with CXCL2 and CXCL3, and IL-1A and IL-6, respectively, indicating that members of CSI participate in cytokine storm. Majority of these TFs are related to inflammatory/immune regulatory processes.

Similarly, during eIF2 signaling/translation, we identified a total of 14 TFs (MXI1, BRCA1, ELF1, SIN3A, E2F4, IRF1, GABPB1, HMGN3, ETS1, SP2, POLR2A, ELK4, CHD2, and CCNT2) expressed in BALF and PBMC (Supplementary Fig. 3, Supplementary Data 4). This set of proteins is involved in the dynamic regulation of eIF2 signaling/translation-related genes over the course of time. Predominantly, we found that 12 master regulators (CHD2, SIN3A, ETS1, MXI1, CCNT2, E2F4, ELK4, IRF1, GABPB1, ELF1, POLR2A, and HMGN3) of eIF2 signaling/translation are involved in dynamic transcriptional regulation at 24-hour post infection. Interestingly, we revealed that 12 TFs also regulated 9 CSPs (AP2M1, NDUFA10, PHB, PPIA, PPP1CA, RPS20, RTN4, SCCPDH, and UBE2I) along with eIF2 signaling/translation-related genes.

Correspondingly, during protein ubiquitination, we identified a total of nine TFs (IRF1, CCNT2, BRCA1, MXI1, CHD2, POLR2A, SIN3A, E2F4, and HEY1) expressed in BALF and PBMC (Supplementary Data 4). Interestingly, in the gene subset related to T cell receptor regulation of apoptosis, we identified a total of 11 TFs (E2F4, ELF1, CCNT2, ETS1, ELK4, HMGN3, SP2, IRF1, GABPB1, MXI1, and SIN3A) expressed in BALF and PBMC that participate in the regulation of this pathway genes including members of CSPs (Supplementary Data 4). Overall, we demonstrated the dynamic transcription patterns of CSI genes and CSPs that participate in cytokine storm, eIF2 signaling/translation, protein ubiquitination pathway and T cell receptor regulation of apoptosis (Supplementary Fig. 3, Supplementary Data 4). Finally, we identified a set of TFs that potentially regulate the above mentioned pathways’ genes at various stages of SARS-CoV-2 infection in COVID-19 patients.

## Discussion

In the last two decades, intra- and inter-species interactomes have been generated in a number of prokaryotes and eukaryotes including human, mouse, worm and plant models^5, 6, 10, 33^. Investigating such interactomes has indicated that diverse cellular networks are governed by universal laws, and led to the discovery of shared and distinct molecular components and signaling pathways implicated in viral pathogenicity. In the present study, we constructed a <u>C</u>alu-3-specific human-<u>S</u>ARS-CoV-2 <u>I</u>nteractome (CSI) by integrating the lung epithelial cells-specific co-expression network with the human interactome. We determined that CSI displayed features of scale-freeness and was enriched in different centrality measures. Identification of structural modules displayed the relationships with a set of functional pathways in CSI. In-depth network analyses revealed 28 most influential nodes. Additional noteworthy findings pertain to SARS-CoV-2 transcriptional signatures, regulatory relationships among diverse pathways in CSI and overall SARS-CoV-2 pathogenesis including the cytokine storm.

We constructed a comprehensive and robust CSI, a human-viral interactome that displayed scale free properties (*r*^2^= 0.91; Fig. 1d). We also showed that the SARS-CoV-2 interacting proteins (SIPs) exhibit increased average centrality indices compared to the remaining proteins in the network (Fig. 1d, Supplementary Fig. 1a, b). Numerous human-viral interactomes have previously been generated to uncover global principles of viral entry, infection and disease progression. These include human T-cell lymphotropic viruses, Epstein-Barr virus, hepatitis C virus, influenza virus, human papillomavirus, dengue virus, Ebola virus, HIV-1, and SARS-CoV^12, 13, 14, 15, 16, 17, 18, 19, 20, 24, 25, 26, 27, 28^, and all of these interactomes exhibited a power law distribution. Another significant tenet of interactomes is the existence of modular structures or modules, defined as sets of densely connected clusters within a network that exhibit heightened connectivity among nodes within a module. Such nodes within a module have previously been deemed to possess similar biological function or belong to the same functional pathways^39^. Since nodes in CSI not only form protein complexes but also coexpress specifically to coronavirus infection, we extracted several functional modules from our network (Fig. 2 e-k). The mostly highly connected module pertains to eIF2 signaling, and is comprised of protein translation-related proteins such as RPS and RPLs. Indeed, these ribosomal proteins have been shown to interact with viral RNA for viral proteins biosynthesis, and are subsequently required for viral replication in the host cells^40^ Noteworthy, two ribosomal proteins, RPL36 and RPS20, found to interact with several SARS-CoV-2 viral factors. Moreover, both of these proteins are also <u>C</u>SI <u>S</u>ignificant <u>P</u>roteins (CSPs) that harbor increased centrality measures (Fig. 3e). Intriguingly, RPS20 has been demonstrated to operate as an immune factor that activates TLR3-mediated antiviral. It remains to be addressed whether RPS20 is a “double whammy” target of SARS-CoV-2 for (1) hijacking this important factor for viral translation and replication, and (2) suppressing a critical immune signaling pathway. Regardless, ribosomal proteins are critical targets of numerous viruses and play equally essential roles in developing antiviral therapeutics^40^

The ubiquitin proteasome system (UPS) constitutes the major protein degradation system of eukaryotic cells that participate in a wide range of cellular processes, and another critical target of diverse viruses^41^. UPS plays an indispensable role in finetuning the regulation of inflammatory responses. For instance, proteasome-mediated activation of NF-κB regulates the expression of proinflammatory cytokines including TNF-α, IL-1β, IL-8. Similarly, UPS is indispensable in the regulation of leukocyte proliferation^41^. The UPS is generally considered a double-edged sword in viral pathogenesis. For example, UPS is a powerhouse that eliminates viral proteins to control viral infection, but at the same time viruses hijack UPS machinery for their propagation^41^. In case of herpes simplex virus type 1, Varicella-zoster virus and Simian varicella virus, induction of NF-κB – mediated host innate immunity is suppressed by the manipulation of UPS components^42^. Moreover, it was revealed that UPS plays crucial roles at multiple stages of coronaviruses’ infection^43^. In our study, the ubiquitin proteasome module was composed of several members of 26S proteasome ATPase or non-ATPase regulatory subunits, which includes two CSPs, PSMD8 and PSMA2 (Fig. 3e). It still needs to be determined whether these two CSPs play important roles in the expression of proinflammatory cytokines and are potentially involved in the cytokine storm. While the mechanistic evaluation of SARS-CoV-2 interaction with these two high-value targets needs to be explored, both the mRNA and protein expression corresponding to PSMD8 was recently shown to be decreased up to 30% in aged keratinocytes^44^. Since reduced proteasome activity results in aggregation of aberrant proteins that perturb cellular functions, we hypothesized that SARS-CoV-2 targets these CSPs to interfere with ER-mediated cellular responses. Another noteworthy module is the T cell receptor regulation of apoptosis. Indeed, it was recently reported that SARS-CoV-2 infection may cause lymphocyte apoptosis demonstrated by overall cell count and transcriptional signatures in PBMC of COVID-19 patients^36, 45^. Another significant CSP in this pathway is MTCH1 (Fig. 3e), a proapoptotic protein that triggers apoptosis independent of BAX and BAK^46^. We hypothesized that cytokines-mediated induction of cytokine storm is partially dependent on the SARS-CoV-2 interaction with MTCH1. Taken together, our module-based functional analyses identified several novel molecular components, structural and functional modules, and overall provided insights into the pathogenesis of SARS-CoV-2.

Our network topology analyses discovered 28 CSI Significant Proteins (CSPs) that have been implicated in several above described modules and pathways (Fig. 3e). To provide a system-wide perspective of the importance of these CSPs in COVID-19, we categorized these CSPs into three groups based on their possible functionality. Group-1 includes CSPs that are potentially relevant to modifying host response following SARS-CoV-2 infection. These include EEF1A1, ETFA, MRPS27, MRPS5, MTCH1, NDUFA10, RAB1A, RAB2A, RAB5C, RAB7A and RHOA (Fig. 3e). We hypothesized that such CSPs are important in creating protective environment in host tissue following the viral infection. For example, RAB and RHO group of ras proteins may be involved in augmenting inflammatory signaling pathways. While antioxidants regulating mitochondrial and cytoplasmic proteins are possibly important in regulating and maintaining redox homeostasis^47^, another CSP, SCCPDH, is involved in the metabolic production of lysine (Lys) and α-ketoglutarate (α-kg)^48^ Intriguingly, L-lysine supplementation appears to be ineffective for prophylaxis or treatment of herpes simplex lesions^49^. We hypothesized that SARS-CoV-2 may target SCCPDH to hijack the biosynthesis of this essential amino acid for its benefit. Group-2 CSPs that we identified are likely to be hijacked by SARS-CoV-2 for its entry, proliferation and survival in the host tissue. In this category, one of the most important CSPs is prohibitin (PHB; Fig. 3e). PHB is an important protein shown to be a receptor for dengue and Chikungunya viruses^50, 51^. Although it has been shown that ACE2 serves as the main receptor for SARS-CoV-2 entry into the cells^52^, it is quite interesting that pathogenesis of the viral infection is not significantly different between the populations of hypertensive patients who receive or don’t receive ACE2 inhibitors^53, 54, 55, 56^. Therefore, it is plausible that under certain physiological conditions when SARS-CoV-2 does not engage with the ACE2 receptor for its entry into the cell, PHB serves as an alternative receptor. Another CSP Integrin β1 encoded by ITGB1 was recently shown to be required for the entry of Rabies Virus^57^ Whether ITGB1 could also promote the entry of SARS-CoV-2 is another question that needs to be addressed. MEPCE is another important enzyme involved in RNA stabilization by capping the 5’ end of RNA with methyl phosphate^58^. It is also likely that MEPCE is utilized by the COVID-19 virus for stabilization of its RNA in the host tissue. Similarly, PPP1CA was shown to regulate HIV-1 transcription by modulating CDK9 phosphorylation^59^, and thus is potentially involved in the gene regulation of SARS-CoV-2. As discussed above, PSMA2 and PSMD8 are the two proteasomal CSPs^42, 43, 44^. While infecting the lung epithelium, SARS-CoV-2 may utilize these UPS proteins for the fusion with the host cell membrane (Fig. 3e). Similarly, NUP98 can also be utilized for viral entry into the nucleus. Additional three CSPs in this category, RPL36 and RPS20^40, 60^ as well as SRP72^61^, could be employed for viral transcription and protein synthesis (Fig. 3e). Finally, Group-3 CSPs are proteins, which SARS-CoV-2 may utilize both to facilitate its proliferation as well as to induce a conducive environment in the host tissue for its sustenance and pathogenesis (Fig. 3e). These CSPs include AP2M1, CSNK2B, EEF1A1, ETFA, LARP1, RTN4 and UBE2I. Among these CSPs, EEF1A1, PPIA, PSMA2, PSMD8, RAB1A, RAB2A, RAB5C, RAB7A, RHOA and UBE2I are identified as the ones that are potentially associated with the pathogenesis of the cytokine storm as observed in some severely affected patient populations. Intriguingly, EEF1A1, a target of several viruses, is known to be activated upon inflammation^62^. This CSP is independently identified as one of the major regulators in human-SARS-CoV-2 predicted interactome^63^. The CSPs, which regulate protein folding and translation, for example EIAF1, could be utilized by SARS-CoV-2 to halt host protein translation, folding and protein quality control. In addition, we also identified E2F4, TBX3 and SMARCB as first neighbors of some of these CSPs. These CSPs complexes play key roles in promoting cell death, causing inflammation and acting enzymatically as viral integrases. Collectively, these CSPs and their first neighbors could directly and indirectly perform intricate pathopysiological functions but those mentioned here could be the key effects of COVID-19 on host tissue dysregulation. This classification is also crucial for the design of effective therapeutic interventions against COVID-19. Finally, we presented transcriptional modeling of CSI genes including CSPs that participate in cytokine storm, eIF2 signaling/translation, protein ubiquitination pathway and T cell receptor regulation of apoptosis. Thus, these signaling pathways and TFs discovered through our analyses could provide important clues about effective drug targets and their combinations that can be administered at different stages of COVID-19.

In conclusion, we generated a human-SARS-CoV-2 interactome, integrated virus-related transcriptome to interactome, discover COVID-19 pertinent structural and functional modules, identify high-value viral targets, and perform dynamic transcriptional modeling. Thus, our integrative network biology-based framework led us uncover the underlying molecular mechanisms and pathways of SARS-CoV-2 pathogenesis.

## Materials and methods

### Human-SARS-CoV-2 interactome data acquisition

To build human interactome, we assembled a comprehensive protein-protein interactions (PPIs) comprising experimentally validated PPIs from STRING database^32^ and four additional proteomes-scale interactome studies *i.e.* Human Interactome I and II, BioPlex, QUBIC, and CoFrac (reviewed in^33^. The resulted human interactome have 18,906 nodes (proteins) with 444,633 edges (interactions). Our human interactome have 1,200 more proteins and 93,189 interactions which was not included in previous study^37^. We collected a total of 394 SARS-CoV-2 interacting proteins (SIPs) from two recent studies encompassing 332 proteins of SARS-CoV-2-human interactions^32^, and 62 proteins of SARS-CoV- and MERS-CoV–human interactions^37^. (Supplementary Data 1).

### Differential gene expression analysis on SARS-CoV, MERS-CoV and SARS-CoV-2 datasets

We obtained microarray data for GSE33267, GSE37827, GSE56677 from GEO database^64^ and used GEO2R, an interactive web tool to generate differential gene expression between infection and mock treatments at their respective time points. Briefly, GEO2R utilizes limma R package. Limma is an R package for the analysis of gene expression microarray data. Specifically, it uses the linear model for analyzing designed experiments and the assessment of differential expression. A threshold of 2 log fold change and FDR ≤ 0.05 was set for differential expression analysis of all microarray experiments. For comparative study of SARS-CoV-2 expression pattern, we downloaded expression data set of RNAs isolated from the bronchoalveolar lavage fluid (BALF) and peripheral blood mononuclear cells (PBMC) of COVID-19 patients^36^. The criteria for filtering out significant genes were kept as adjusted p-value < 0.05 and foldchange > 2

### Generation of Calu-3 Cells-specific Co-expression Networks in response to SARS-CoV and MERS-CoV infection

We mined Calu-3 Cells-specific datasets from GEO database^64^, and downloaded GSE33267 (Wild type), GSE37827 (icSARSCoV) and GSE56677(LoCov). We performed individual weighted gene co-expression network analysis (WGCNA)^65^ package (R version 3.6.1), and constructed three co-expression networks. Moreover, we also generated topological overlap measure (TOM) plots to compute a numerical entity that reflects interconnectedness among genes within a co-expression network. A cut-off of 0.75 was used to export the networks. Subsequently, we merged these networks to generate a comprehensive Calu-3 cells-specific co-expression to study the network connectivity pattern of interactome.

### Network Integration and Topology Analysis

To extract the Calu-3-specific human-SARS-CoV-2 Interactome (CSI) we integrated the merged transcriptomics co-expression network (22, 445 nodes with 10,649,854 edges) and SARS-CoV-2-Human Interactions (12,852 nodes with 84,100 edges) including 373 SIPs. The resulted CSI network has 4,123 nodes with 14,650 edges including 214 SIPs with all possible interactions including their first and second neighbors (Supplementary 66 Data 1). Network topology analyses was performed using NetworkX^66^ (version 2.4) Python (version 3.7.6) package was used except weighted *k*-shell-decomposition for which we downloaded w*k*-shell-decomposition Cystoscope App (version 1.0). Cytoscape (Version 7.3.2) was used to visualize all the networks.

### Gene Ontology Functional Enrichment Analysis

The functional enrichment analysis was done by Kyoto Encyclopedia of Genes and Genomes (KEGG), ingenuity pathway analysis (IPA), WikiPathways, GO biological process, ClueGO, and enricher for human phenotype ontology and rare diseases term with their statistically significant parameters^67^.

### Reconstructing SARS-CoV Responsive Dynamic Regulatory Events

Interactive visualization of dynamic regulatory networks (iDREM) is a method which incorporates static and time series expression data to reconstruct condition-specific reaction network in an unsupervised manner^28^. Additionally, the regulatory model identifies specific stimulated pathways and genes, which uses statistical analysis to recognize TFs that vary in activity among models. We implemented iDREM on 4,952 cumulative differentially expressed genes across 72 hours of SARS-CoV infection with log2 normalization for dynamic regulatory event mining with all human 954,377 TFs/targets collections from encode database^68^. The dynamic activated pathways regulated by TFs was generated by EBI human gene ontology function.

### Statistical analyses

Hypergeometric test, linear regression (*r*^2^), and Student *t*-test were performed using R version 3.3.1 as well as online Stat Trek tool.

## Data Availability

All datasets used for this study are accessible through Supplementary Data files.

## Acknowledgements

This work was supported by the National Science Foundation (IOS-1557796) to M.S.M., and U54 ES 030246 from NIH/NIEHS to M. A. Thanks to Dr. K. Mukhtar for critical reading the manuscript.

## Contributions

M.S.M. N.K. and M.A. conceived the project. N.K. B.M. and A. M. performed networkbased and statistical analyses. M.S.M. and M.A. wrote the first draft of the manuscript except methods. N.K. and B.M wrote the methods section. All the authors discussed the results and critically reviewed the manuscript and provided valuable comments/edits.

## Ethics declarations

### Electronic supplementary material

Supplementary Information

Description of Additional Supplementary Files

Supplementary Data 1

Supplementary Data 2

Supplementary Data 3

Supplementary Data 4

