## Supplementary Information for "Integrative Network Biology Framework Elucidates Molecular Mechanisms of SARS-CoV-2 Pathogenesis"

### Integrative Network Biology Framework Elucidates Molecular Mechanisms of SARS-CoV-2 Pathogenesis and Potential Therapeutic Intervention

Nilesh Kumar[1], Bharat Mishra[1], Adeel Mehmood[1, 2], Mohammad Athar[3]* & M. Shahid Mukhtar[1, 4, 5]*

Affiliations

[1] Department of Biology, 464 Campbell Hall, 1300 University Boulevard, University of Alabama at Birmingham, Alabama 35294, USA

[2] Department of Computer Science, University of Alabama at Birmingham, 1402 10th Ave. S. , Birmingham, AL 35294, USA.

[3] Department of Dermatology, School of Medicine, University of Alabama at Birmingham, Alabama 35294, USA

[4] Nutrition Obesity Research Center, 1675 University Blvd, University of Alabama at Birmingham, Birmingham, AL 35294, USA.

[5] Department of Surgery, 1808 7th Ave S, University of Alabama at Birmingham, Birmingham, AL 35294, USA

***Correspondence**

These authors contributed equally: Nilesh Kumar and Bharat Mishra

**Supplementary Text and Figures**

Supplementary information for Supplemental files

Supplementary Data 1: Integrated Calu-3-specific human-SARS-CoV-2 Interactome (CSI)

1. List of SARS-CoV-2 Target Proteins
2. List of interaction of CSI
3. List of CSI Nodes and their Property

Supplementary Data 2: Integrated Calu-3-specific human-SARS-CoV-2 Interactome (CSI) Network Properties

1. CSI Network Analysis Nodes Properties
2. Ingenuity Pathway Analysis (IPA) of CSI MCODE clusters
3. Ingenuity Pathway Analysis (IPA)of CSI Nodes
4. KEGG Pathway Analysis of CSI Nodes

Supplementary Data 3: 28 CSI Significant Proteins (CSPs)

1. 28 CSI Significant Proteins (CSPs) Properties
2. Significant CSPs in seven indices
3. Rare Disease Pathways of CSPs
4. WikiPathways of CSPs

Supplementary Data 4: Dynamic gene regulation modeling of core transcriptional circuitry in SARS-CoV-2 infection by iDREM

1. 228 DEGs in COVID-19 patients derived from bronchoalveolar lavage fluid (BALF) and CSI
2. 215 DEGs in COVID-19 patients derived from peripheral blood mononuclear cells (PBMC) and CSI
3. 4,952 DEGs log fold change expression in SARS-COV
4. 108 iDREM Significant TFs
5. Significant TFs at 24 Hours
6. TFs in eIF signaling/ Translation
7. TFs in Protein Ubiquitination Pathway
8. TFs in T cell receptor regulation of apoptosis
9. TFs in Cytokine Storm
10. TFs-Targets enriched in Cytokine Storm during iDREM

### Supplementary Figures

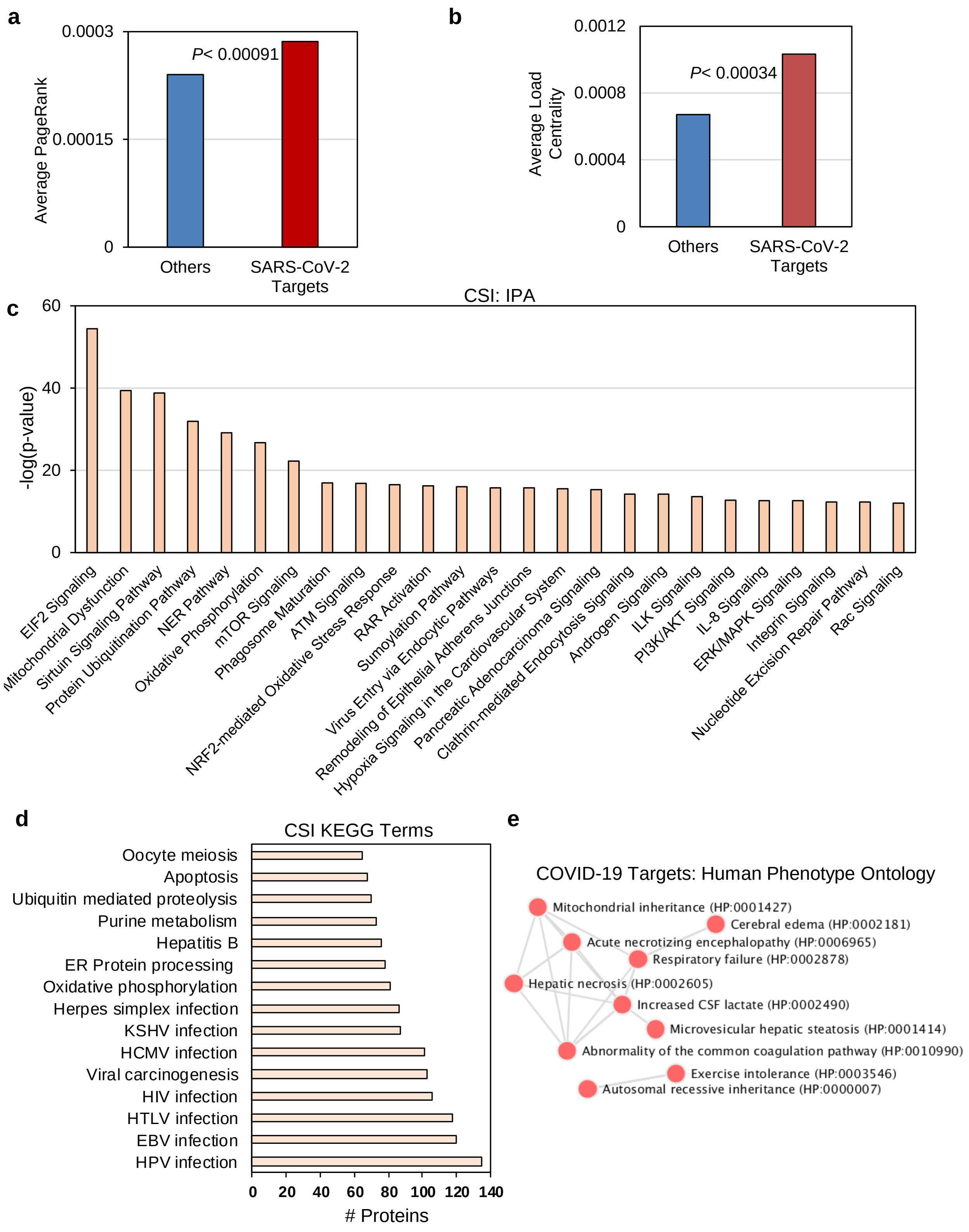

**Supplementary Figure 1: SARS-CoV-2 Interacting Proteins (SIPs) additional structural and functional properties are enriched in Calu-3-specific human-SARS-CoV-2 Interactome (CSI).**

**a** Average PageRank of SIPs (0.000286) is significantly higher than that other interacting proteins (0.00024) in CSI network (*t*- test, $P < 0.00091$). **b** SIPs display significantly increased average load centrality of (0.0011) compared to other interacting proteins (0.000672) in CSI network (*t*- test, $P < 0.00034$). **c** Ingenuity Pathway Analysis (IPA) identified significantly enriched canonical pathways in CSI proteins (-log(*P*-value) ≥12). **d** Kyoto Encyclopedia of Genes and Genomes (KEGG) pathway analysis identified infection to viruses, oxidative phosphorylation, ER protein processing and apoptosis that are significantly enriched in biochemical and metabolic pathways in CSI proteins (*P*-value ≤0.05). **e** Significantly enriched human phenotype ontology is identified using Enrichr. Mitochondrial inheritance, hepatic necrosis, respiratory failure and abnormality of the common coagulation pathway terms are enriched.

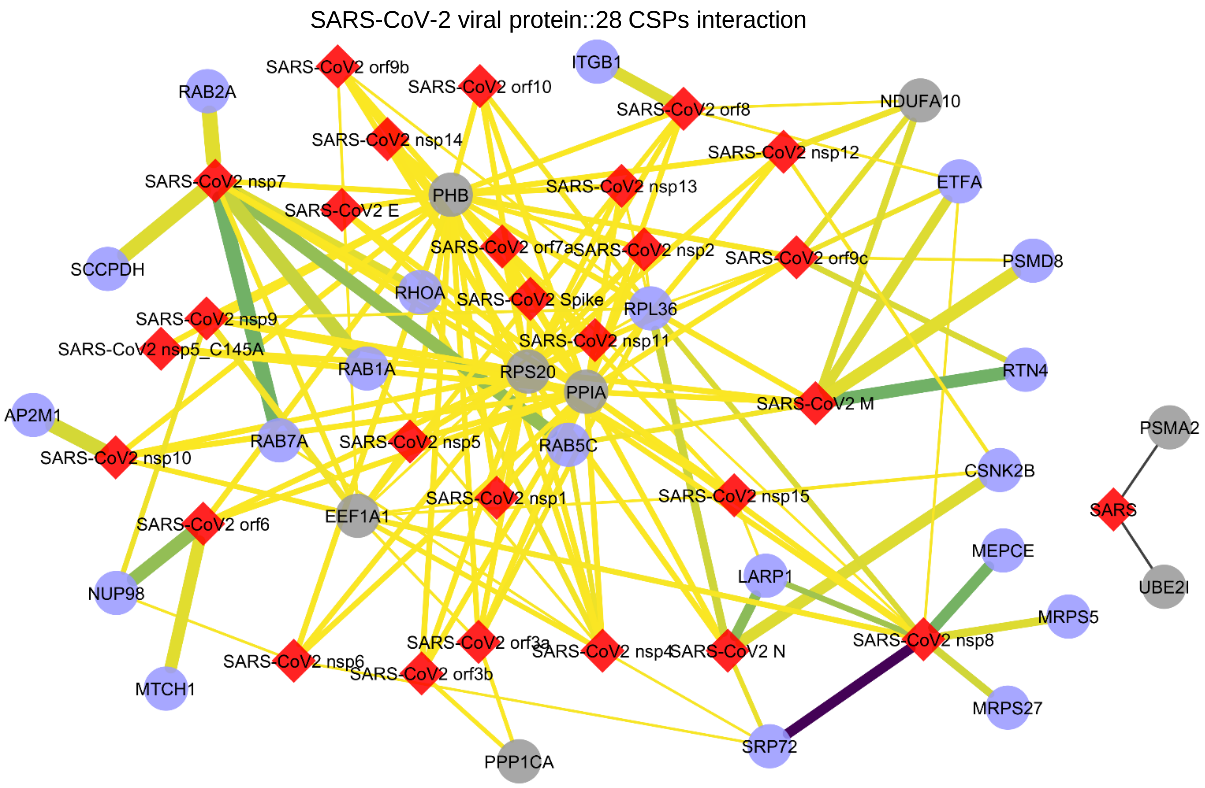


**Supplementary Figure 2: <u>C</u>SI <u>S</u>ignificant <u>P</u>roteins (CSPs) interacting SARS-CoV-2 proteins**

Network representation of SARS-CoV-2 viral protein interaction with 28 CSPs (Nodes: Red= viral proteins, Blue= CSPs significantly targeted by viral protein, grey= CSPs with insignificant viral protein interaction; edge width= MIST score, edge color= AvgSpec).

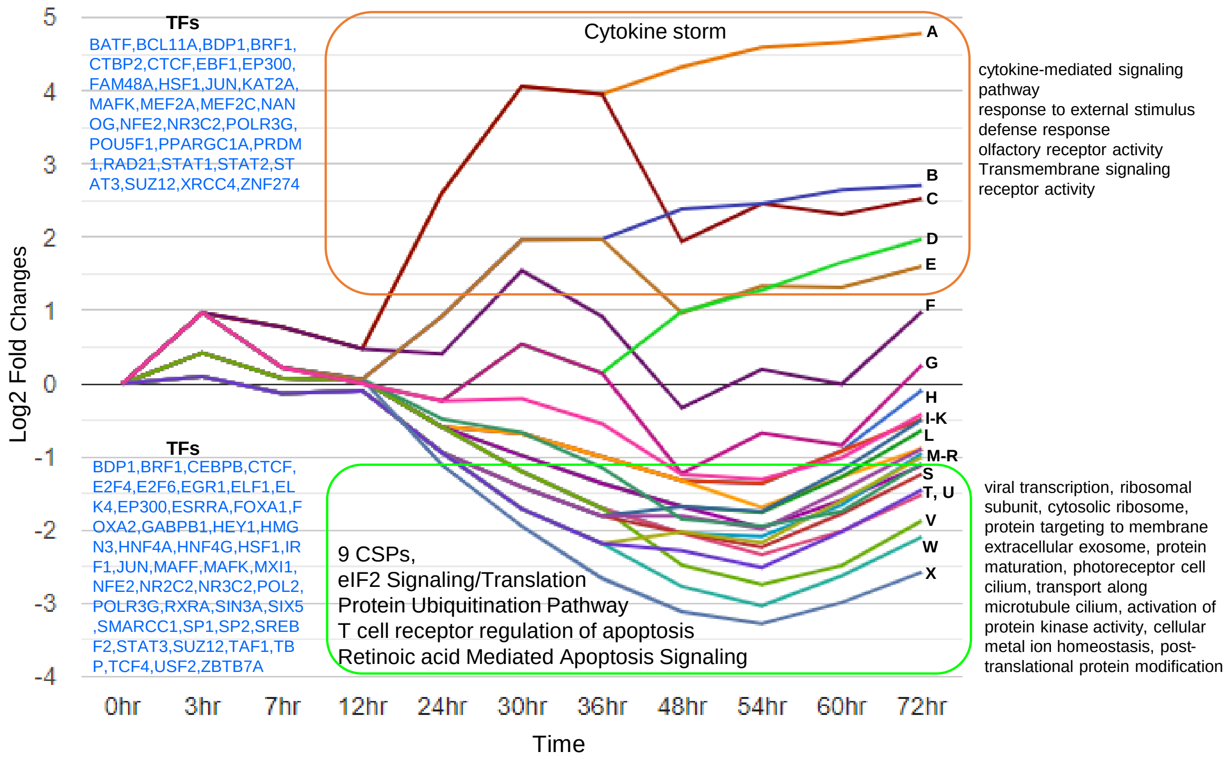


**Supplementary Figure 3: 26 significant path expression pattern of 4,952 differentially expressed genes (DEGs) in SARS-CoV infection across 72 hours (GSE33267).**

Dynamic regulatory event mining of 4,952 cumulative DEGs in SARS-CoV across 72 hours of infection with log2 fold change expression. The regulators only expressed in BALF and PBMC transcriptomes are highlighted. Total significant bifurcated paths (A-X) were identified based on respective significant regulators (TFs) (*P*< 0.05). Top five path expressions (A-E) are mostly enriched by cytokine storm genes (*P*< 0.05). Additionally, these five paths are enriched by cytokine-mediated signaling pathway, response to external stimulus, defense response, olfactory receptor activity, and transmembrane signaling receptor activity. Most significant regulators of cytokine storm genes are (BATF, BCL11A, JUN, MEF2A, NFE2, RAD21, STAT1, STAT2, STAT3, and SUZ12). While, bottom six path expressions (S-X) are mostly enriched by 9 CSI significant proteins, eIF2 Signaling/Translation, protein ubiquitination pathway, T cell receptor regulation of apoptosis, and retinoic acid mediated apoptosis signaling ($P < 0.05$). Additionally, these six paths are enriched in viral transcription, ribosomal subunit, cytosolic ribosome, protein targeting to membrane extracellular exosome, protein maturation, photoreceptor cell cilium, transport along microtubule cilium, activation of protein kinase activity, cellular metal ion homeostasis, post-translational protein modification. There are 43 significant TFs enriched in these processes.
